# Defensive freezing sharpens threat-reward information processing during approach-avoidance decision making

**DOI:** 10.1101/2024.08.29.610250

**Authors:** Felix H. Klaassen, Bob Bramson, Jan-Mathijs Schoffelen, Lycia D. de Voogd, Karin Roelofs

## Abstract

People regularly face approach-avoidance dilemmas which require minimization of potential threat whilst maximizing potential reward. Defensive reactions to threat, such as transient states of freezing, influence integration of reward/threat information in the dorsal Anterior Cingulate Cortex (dACC). However, the mechanism of this integration between internal state and external value (state-value integration) remains unknown. Here, we decoded approach-avoidance decisions under threat using high-precision magnetoencephalography (MEG). Threat-induced cardiac deceleration (indicative of defensive freezing) was trial-by-trial associated with more pronounced effects of reward and threat magnitudes on approach-avoidance choices. Time-resolved decoding of threat-reward information from neural signals predicted approach-avoidance choices several seconds before the actual response. Crucially, during freezing, 6-12 Hz coherence between threat-reward information and dACC neural activity increased, suggesting that defensive freezing sharpens threat/reward processing during approach-avoidance decision making. These findings provide a potential neural mechanism by which threat-induced freezing can facilitate information integration, essential for optimal decision making under threat.

## Introduction

We are regularly faced with approach-avoidance conflict situations that require the minimization of danger whilst maximizing potential benefits^1–3^. Whilst avoiding threatening situations might prevent physical or psychological harm, it may come at the cost of missing potential opportunities. Approach-avoidance decisions are often modelled within value-based computational frameworks that describe how reward and threat values influence behavior^4^. However, biological agents such as humans have not developed to make decisions based solely on objective reward and threat values^5^. Rather, and especially in threatening situations, the current psychophysiological state of our bodies may affect what information we seek out, and how that information influences our decisions^6–8^. Here, we show how defensive freezing states during approach-avoidance decision making, assessed via cardiac deceleration, sharpen the influence of threat and reward prospects on behavior and human brain activity.

The freezing response is a parasympathetically-driven threat-anticipatory reaction characterized by bodily immobility and heart rate deceleration^9–11^. Recent models detail that during freezing cholinergic projections from the midbrain and forebrain influence neural processing throughout the cerebral cortex, sharpening visual perception, action preparation, and potentially also value-based processing^12–19^. Accordingly, we confirmed the involvement of the amygdala in computing threat value and of the dACC in threat-reward value comparison as a function of defensive freezing states^8^. However, how the brain performs this state-value integration process during decision making, and how those operations change during freezing states, remains unclear.

During decision making, the evaluation of conflicting potential reward and threat outcomes can follow a sequential neural representation pattern in prefrontal cortex^20,21^. Such a pattern might reflect competition through mutual inhibition between neural ensembles, informed by threat and reward information from connected regions^22,23^. Here we consider the possibility that, in order to arbitrate approach-avoidance decisions, the brain might iteratively represent the potential reward and threat outcomes, resulting in rhythmic magnetoencephalographic (MEG) patterns of reward and threat information over time, until a decision is reached^22^. Interestingly, in rodents, the organization of defensive responses such as freezing has been associated with 4 – 12 Hz oscillatory activity across frontal cortico-limbic structures also involved in value-based choice^24^. Based on these findings, we hypothesize that during decision making, neural threat-reward information follows a rhythmic representation pattern, and that altered threat and reward processing during freezing is supported by changes in the low-frequency rhythmic dynamics of these neural representations in the dACC.

We tested these hypotheses by capitalizing on the power of a within-subjects design and trial-by-trial analyses. Nine human participants each performed 5 sessions of an approach-avoidance conflict decision-making task^8^ in which they chose to approach or avoid targets associated with varying reward and threat magnitudes (0-5 euros/electrical shocks; **Figure 1a**). To maximize within-subjects reliability, we recorded neural activation using high-precision MEG with individualized head casts. We find that defensive freezing, assessed through trial-by-trial cardiac deceleration (see e.g., refs^11,14,17,25,26^), sharpens the effect of threat and reward on approach-avoidance choices. Using classification analysis, we reveal that neural threat-reward information decoded from the MEG signal predicts the actual approach-avoidance responses made seconds later, and is synchronized with low-frequency rhythmic activity in the dACC (6 – 12 Hz). Moreover, this correlation between low-frequency activity and the strength of the decoded threat-reward representation is more pronounced during stronger cardiac-indexed freezing.

**Figure 1.**
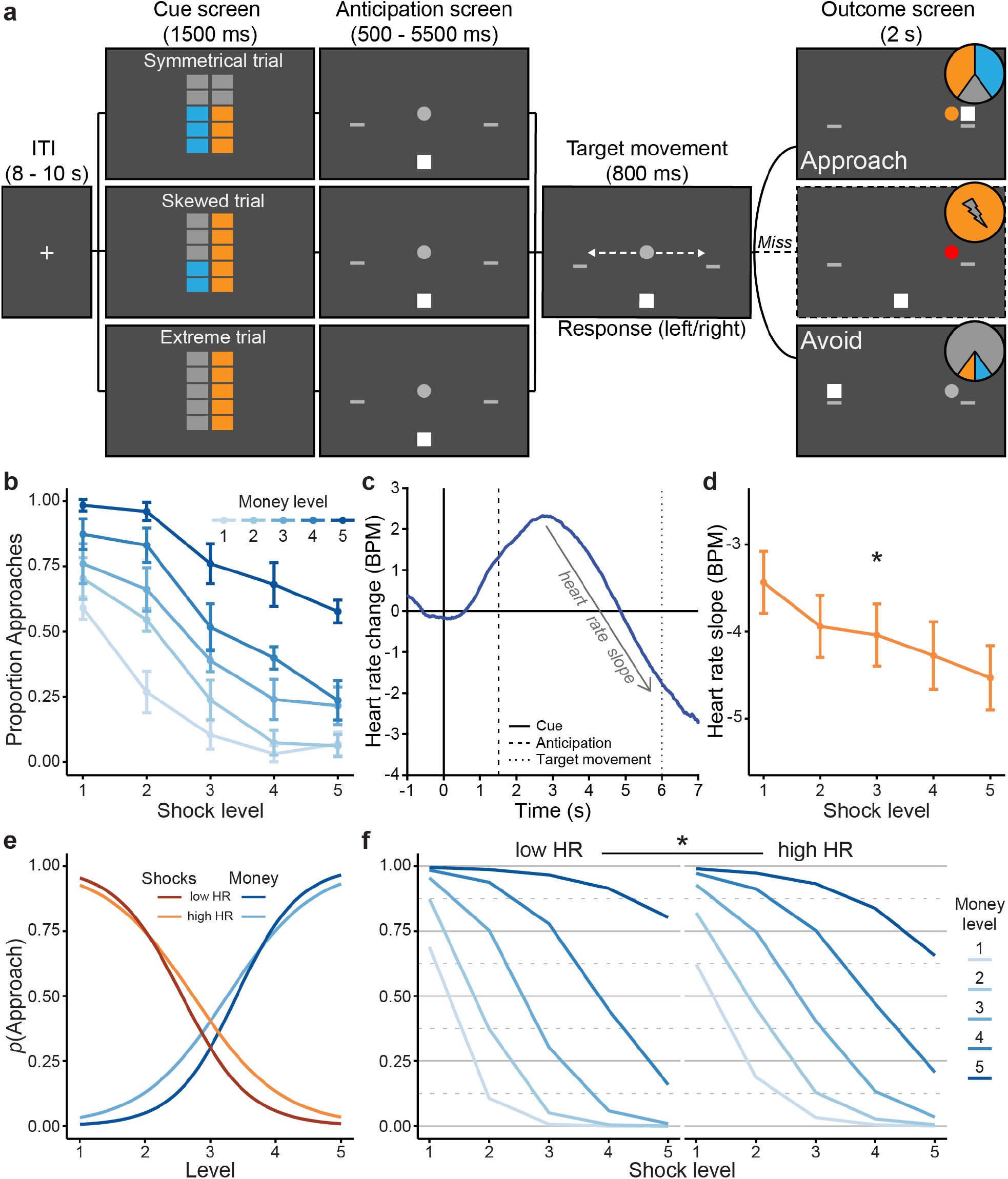
a) Approach-avoidance conflict task. (**a**) On each trial, participants approached or avoided a target associated with varying reward and threat magnitudes (0-5 euros/shocks). The reward and threat magnitudes on offer were indicated during the cue screen at the start of the trial by two stacks of orange and blue boxes. There were three classes of stimuli: symmetrical trials in which the magnitudes of both outcomes were identical, skewed trials in which the magnitudes were different (but higher than 0), and extreme trials in which the magnitude for one outcome was 5 and for the other was 0. Next, during the anticipation screen, the participant was indicated by a white square at the bottom of the screen, while the target was indicated by a grey circle (presented centrally). After a variable interval, the target started to move either to the left or right, initiating the response window. To approach the target, participants had to jump to the same location as the target, whereas to avoid it participants had to jump to the opposite direction (using buttons under their left and right index fingers). Approach was associated with a high probability to receive either the money (40%) or shocks (40%), and a small probability of no outcome (20%). Avoidance was associated with small probabilities of money (10%) and shock (10%) outcomes, and a large probability of no outcome (80%). Missed responses always led to shocks. Participants were informed on these contingencies. The selected outcome (money/shocks/nothing) was presented during the outcome screen by color coding of the target (blue/orange/gray). Additionally, shock outcomes were accompanied by administration of the indicated number of electrical shocks, and money outcomes were accompanied by presentation of a cross (‘+’) centered in the target (the number of flashes corresponding to the money level). Orange/blue color assignment for reward/threat magnitudes was counterbalanced between participants. Dashed arrows were not present in the actual task. (**b**) Increased money and shock levels were associated with more approach and avoidance choices, respectively. (**c**) We observed significant heart rate (HR) deceleration over time (i.e., a negative slope) during the anticipation window, indicating a psychophysiological state of anticipatory defensive freezing. (**d**) Cardiac deceleration was more pronounced for higher shock levels. (**e**) The effects of money and shock levels on choice were more pronounced in trials with stronger cardiac deceleration. (**f**) A three-way interaction between money, shocks, and heart rate indicated that the interaction between money and shocks was stronger in trials with more cardiac deceleration. All effects were tested on a trial-by-trial level, low and high HR conditions (indicating steep vs. shallow negative slopes) were only created for plotting purposes. Error bars indicate 1 standard error of the mean (SEM), asterisks indicate significant effects (i.e., HDI_95%_ does not include 0). Solid, dashed, and dotted vertical lines in (**c**) reflect cue, anticipation, and target movement onsets, respectively. Note that target movement onset was uniformly jittered from 6 – 7 s relative to the cue onset. **e)** and **f)** are conditional-effect plots of the probability to approach ‘p(approach)’ extracted from the Bayesian mixed-effects models (BMMs).

## Results

Participants performed an approach-avoidance decision-making task in which they could win money (0-5 euro) at the risk of receiving electrical shocks (0-5 shocks each trial; **Figure 1a**). Each offer was presented for 1.5 seconds, 5 to 6 seconds prior to the response window. This delay is needed to allow assessment of the slowly developing cardiac deceleration across time^8,17^. Extreme trials, where only threat or reward was on offer, were used to train a classifier to distinguish reward from threat information in the neural signal. This classifier was then applied to independent trials with a combination of both threat and reward on offer, providing a moment-by-moment estimate of the extent to which the neural signal resembled threat or reward information. We used this decoded signal to predict subsequent approach-avoidance choices, and show how neural threat-reward processing changed during defensive freezing states (indicated by cardiac deceleration).

### Defensive freezing is associated with stronger influence of threat and reward on decision making

Previous work has repeatedly shown that approach-avoidance choices are influenced by both potential threat and reward, and moderated by psychophysiological freezing states which can be indexed through cardiac deceleration^8,11,17,25^. Here, we confirm those effects, showing that participants approached more for higher money levels and avoided more for higher shock levels (with a marginally significant interaction; B_money_ = 2.56, HDI_95%_ = [1.43, 3.78]; B_shocks_ = −2.42, HDI_95%_ = [−3.42, −1.39]; B_money:shocks_ = 0.72, HDI_90%_ = [0.03, 1.37]; **Figure 1b**). During the anticipation window, heart rate decelerated (B_intercept_ = −3.99, HDI_95%_ = [−5.75, −2.04], **Figure 1c**), and significantly more so for higher shock levels, indicative of defensive freezing (B_shocks_ = −0.28, HDI_95%_ = [−0.5, −0.07], **Figure 1d**). There was only a marginal effect of money level on cardiac deceleration (B_money_ = −0.18, HDI_90%_ = [−0.34, −0.01]).

In line with our previous work, approach-avoidance decisions depended on trial-by-trial differences in defensive freezing. First, the effects of money and shock levels on choice were more pronounced for stronger cardiac deceleration (B_money:HR_ = −0.41, HDI_95%_ = [−0.8, −0.07]; B_shocks:HR_ = 0.33, HDI_95%_ = [0.07, 0.61], **Figure 1e**). Second, a significant three-way interaction indicated that the interaction between money and shock levels on choice was stronger for more negative slopes (i.e., cardiac deceleration), compared to more positive heart rate slopes (B_money:shocks:HR_ = −0.24, HDI_95%_ = [−0.47, −0.02], **Figure 1f**). Cardiac deceleration was also associated with faster response times in general (B_HR_ = 0.02, HDI_95%_ = [0.003, 0.04]). Together, these results confirm that the integration of potential threat and reward outcomes into a decision depends on the current defensive cardiac state. Next, we addressed our main research question when and how the brain computes this state-value integration.

### Threat and reward information decoded from the neural signal predicts approach-avoidance choices

In order to test how defensive freezing might change the influence of threat and reward information on decisions, we acquired high-precision MEG data from our participants that each performed our task for 5 hours in total, using individualized head casts. We trained an individualized classifier to dissociate extreme threat (5 shocks, 0 euro) from extreme reward trials (5 euro, 0 shocks). We reasoned that training a classifier on the extreme trials would yield the cleanest estimate of neural threat and reward information_20_. The resulting classifier performed significantly above chance on left-out extreme trials (average classification accuracy of 69.8%, range 61.6 - 76.5, chance being 50%; *t*(8) = 13.8, *p* < .0001, **Figure 2a**) and peaked for all participants during the cue screen. Classification accuracy could not be explained by eye movements (see **Supplementary Figure S1**) or cardiac differences between conditions that manifest later in the trial. For time-by-time performance of the classifier, see **Supplementary Figure S2**.

**Figure 2.**
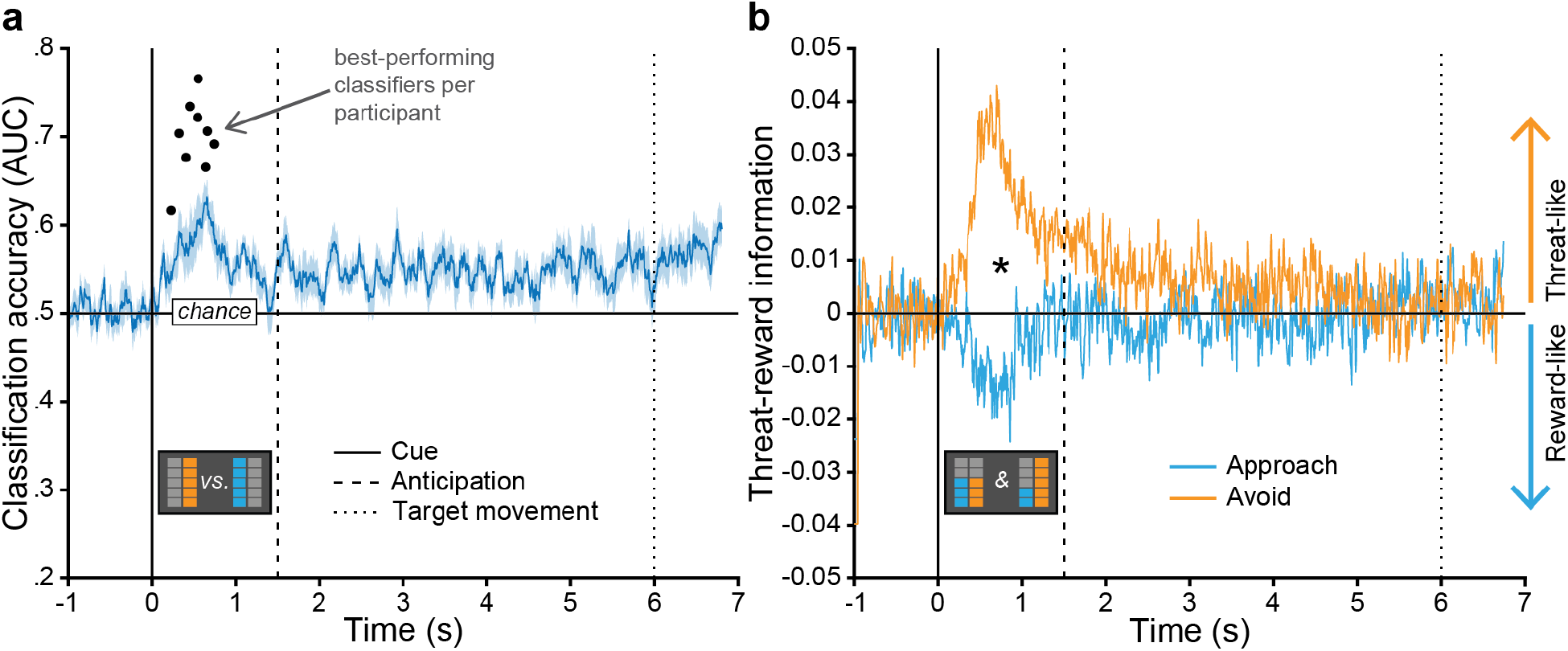
Threat and reward information decoded from the neural signal predicts approach-avoidance choices. (**a**) Classification of extreme reward vs. extreme threat trials across participants was significantly above chance (50%) during the cue screen (0 – 1.5 sec) and peaked between 233 – 746 ms. Black dots indicate individual participant classifiers, average classification accuracy across participants per sample indicated in blue, shaded area reflects +/-1 SEM. (**b**) Cross-classified trial-by-trial threat-reward information (TR values) in symmetrical and skewed trials was predictive of subsequent approach-avoidance choices. More positive TR values reflect more threat-like information, whereas more negative TR values reflect more reward-like information. Solid, dashed, and dotted vertical lines reflect onset of the cue, anticipation, and target movement screens, respectively (note that the target movement onset was uniformly jittered from 6 – 7 s relative to the cue onset).

We then selected the best-performing classifier per participant (black dots in **Figure 2a**) and applied these to each timepoint of the independent symmetrical and skewed trials containing *both* threat and reward information. The resulting timeseries of threat-reward values (**TR values**) reflects to what extent neural information at each timepoint is more threat-like or more reward-like. Crucially, these TR values predicted subsequent approach-avoidance choices made seconds later during the target movement screen. Namely, participants approached more often in trials where the neural signal during cue presentation more strongly resembled extreme reward, whereas trials containing more threat-like information resulted in more avoidance choices (**Figure 2b**; B_TRval_ = −0.20, HDI_95%_ = [−0.34, −0.06]). Importantly, this effect remained significant when only considering the symmetrical trials in which the objective threat and reward magnitudes were identical (B_TRval_symm_ = −0.21, HDI_95%_ = [−0.38, −0.04]), demonstrating that this finding is not driven by potential biases in decoding due to imbalances in onscreen visual information but rather by the meaning conveyed by the stimulus.

### Neural threat and reward representations are rhythmic and more strongly synchronized to dACC during defensive freezing

Now that we computed from the neural signal a behaviorally relevant time-resolved estimate of threat-reward information processing, we asked how this threat and reward information processing is changed during defensive freezing. For this we focused on the dorsal anterior cingulate cortex (dACC), a region we have shown to be involved in the integration of threat/reward values and freezing states^8^.

Because previous work has shown rhythmic iterations of choice options in prefrontal cortex^20^ we performed a Fourier transform on the TR value timeseries extracted from the non-extreme trials. This revealed rhythmicity in the TR value timeseries, which was particularly pronounced around the 6 – 12 Hz frequency band (**Figure 3a**). We then used a jack-knife procedure^27^ to compute trial-by-trial instantaneous coherence of threat-reward information within this frequency range with neural activity timeseries extracted from the left and right dACC (**Figure 3b)**. Jack-knifing computes coherence iteratively, each time leaving out one trial. The resulting distribution of coherence values across trials reflects the influence of the left-out trial on the average coherence estimate. Note that this procedure flips the direction of the coherence metrics, such that lower values reflect stronger coherence (**Figure 3c**). Trial-by-trial coherence between TR values and dACC rhythmic neural activity was significantly stronger in trials with stronger cardiac deceleration (stronger freezing) in bilateral dACC; dACC_left_: B_HR_ = 0.034, HDI_95%_ = [0.002, 0.068]; dACC_right_: B_HR_ = 0.036, HDI_95%_ = [0.006, 0.06], **Figure 3c**. In other words, in states of stronger cardiac deceleration, activity in the dACC was more strongly associated with 6 – 12 Hz rhythmicity in threat-reward information. Repeating this analysis in control regions showed that this relationship is anatomically specific to dACC and neighboring supplemental motor area (SMA), and not the dorsolateral prefrontal cortex or posterior cingulate cortex (dlPFC, PCC; see **Supplementary Figure S3**). This increase in trial-by-trial 6 – 12 Hz coherence was also marginally associated with more subsequent avoidance responses, indicating its behavioral relevance (dACC_right_: B = 0.078, HDI_90%_ = [0.009, 0.15]; dACC_left_: B = −0.01, HDI_90%_ = [−0.09, 0.07]).

**Figure 3.**
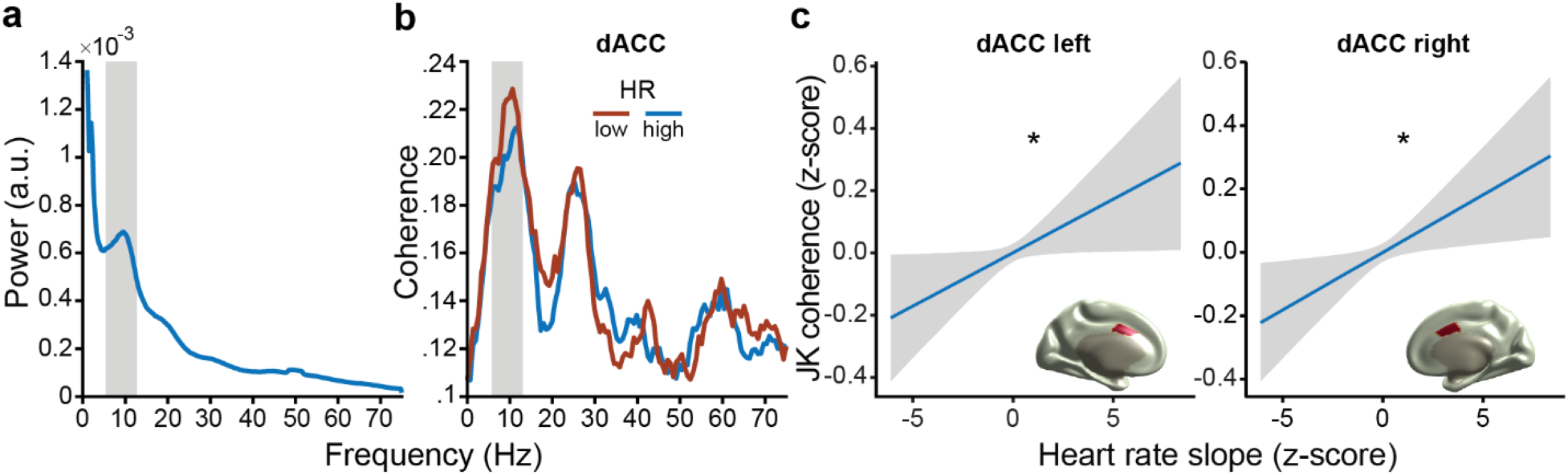
The temporal dynamics of decoded threat-reward information and its moderation by cardiac states. (**a**) Trial-by-trial threat-reward information contains rhythmicity that was particularly pronounced in the 6-12 Hz frequency range. (**b**) Coherence spectrum for reconstructed bilateral dorsal anterior cingulate cortex (dACC) activity with decoded threat-reward information (averaged across hemispheres). For display purposes only, coherence is plotted separately for trials with 25% weakest (blue) vs. 25% strongest (red) cardiac deceleration (i.e., both bins contain equal numbers of trials). Lower HR values indicate stronger cardiac deceleration. (**c**) Trial-by-trial jack-knife (JK) coherence in the left and right dACC with decoded threat-reward information in the 6-12 Hz range (indicated with gray shaded rectangles in **a** and **b**) is stronger during states of cardiac deceleration. Note that the y-axis is flipped; lower values indicate stronger coherence; similarly, lower heart rate slope values on the x-axis indicate stronger cardiac deceleration. Conditional-effect plots in **c** were extracted from the multivariate BMM. Anatomical regions from which MEG signal was reconstructed are indicated in red. Asterisks indicate significant effects (i.e., HDI_95%_ did not include 0).

## Discussion

In this study we test the role of defensive freezing states in decision making under threat and highlight the relation between cardiac deceleration and threat-reward integration. There are three main findings. First, defensive cardiac states moderate reward and threat effects on approach-avoidance decisions. Specifically, stronger cardiac deceleration (indicative of defensive freezing) is related to more pronounced effects of reward, threat, and their interaction on choice, suggesting an upregulation of value-based processing during freezing. Second, neural threat-reward information decoded from whole-brain MEG within the first 600 ms after stimulus presentation predicts actual approach-avoidance responses made seconds later. Third, 6 – 12 Hz rhythmic synchronization between decoded threat-reward signals and neural activity in the dACC is increased during states of cardiac deceleration. This suggests that modulation of low-frequency oscillatory activity in dACC during defensive freezing might support the sharpened influence of reward and threat information on choice behavior. Together, these findings advance our insights into neural dynamics that support decision making under threat. They provide evidence that human defensive psychophysiological freezing states are associated with potentiated neural processing of reward and threat prospects early in the decision-making process, subsequently affecting actions.

Trial-by-trial cardiac deceleration was associated with more pronounced effects of reward and threat magnitudes and their interaction on subsequent approach-avoidance choices. This aligns with reports showing that freezing states are associated with sharpened sensory processing when decisions have to be made based on ambiguous stimuli^16,18,28^, potentially reflecting a role of defensive freezing in reducing decision noise^7,18^. How can this be implemented? During value-based decision making, neural ensembles in frontal cortex alternate representations of potential choice options^20^. Competition between those neural ensembles through mutual inhibition determines which option is chosen^22^, where the balance of excitation/inhibition, combined with the amount of evidence, defines the speed by which the system settles on a decision^29,30^. Speculatively, cholinergic projections from the forebrain to ACC that increase during freezing^19^ might change NMDA receptor functioning in excitatory or inhibitory ACC neurons, thereby changing the gain by which those neural populations drift towards one decision option. Similar processes have been demonstrated for other neuromodulators^31^. The speeding of responses we observe during freezing, independent of reward/threat value, resembles speeding of decisions that occurs with increased overall value or increased value difference between options^32^ and suggests that acetylcholine release might potentiate excitatory neurons, thereby increasing the ramping-to-threshold in dACC circuits. Future work could test this prediction by altering acetylcholine through pharmacological interventions combined with modelling of neural computations in dACC (e.g., see ref^31^).

Threat and reward value decoded early during cue presentation predicts approach-avoidance responses made several seconds later. The fact that the prediction accuracy diminishes over time fits the notion that neural response typically show the strongest transients when competition between options is being resolved^32^, and settle in a low-energy state afterwards^33,34^. These latter states could differ qualitatively from the patterns our classifier was trained on preventing generalization across time^35,36^. The pattern we observe when performing time-by-time classification supports such an interpretation (see **Supplementary Figure S2)**. Namely, a strongly predictive transient during the cue window that does not transfer to later timepoints, and a sustained, although less strongly predicted pattern after the cue disappears.

In a previous fMRI study we showed that dACC is important for integrating external threat and reward information with bodily states when computing approach-avoidance decisions under threat^8^. Here, we extend that knowledge by showing that this integration is associated with increased synchronization of dACC neural populations in less than a second. These oscillatory signatures can be a natural consequence of mutual inhibition, and are often observed during value-based choice^20,22^. Crucially, the slow oscillations involved in these processes might open periods of increased excitability in which long-range projections from other regions can contribute to the computations in dACC^37^. Previous work in non-human primates has shown that neural activation in ACC ramps up and down following reward computations in orbitofrontal cortex when multiple potential reward values are compared^23^. In cases where potential threats have to be taken into account, amygdala projections to dACC could weigh in on these computations^8,38^. Indeed, during freezing, dACC activation is strongly entrained with amygdala activation^24^. When approach actions have to be made in the presence of high conflict, dACC neurons can override threat and bias to approach through projections to the striatum^39^, an area we previously also observed to be involved during this task^8^. Here, we tentatively find that low-frequency coherence of dACC with threat-reward information is related to subsequent avoidance responses, indeed suggesting that this area integrates reward and threat values into a decision. How dACC then translates this decision into an action, and whether freezing states affect this process, remains to be tested^15^.

Our experimental design maximized intra-individual sensitivity by testing few participants for many more repetitions, using individualized head casts^40^. Additionally, our design allowed us to minimize visual differences between reward and threat stimuli, counterbalance their presented location so as to obscure the required left/right response option, and train threat-reward decoders on *anticipatory* threat/reward-related MEG activity in independent trials (i.e., extreme reward/threat trials) whilst testing on trials with the same objective amount of threat/reward information. This alleviates some of the difficulties in previous decoding studies on approach-avoidance choices that had to train on the neural response to either the outcome (e.g., administration of electrical shocks^21^) or on visually highly discriminable stimuli^41^, limiting their sensitivity to anticipatory processing of threat/reward value.

It is unlikely that our classification accuracy can be explained by sources of noise such as eye movements or remaining heartbeat signals in the neural data. Although the accurate prediction of approach-avoidance choices from the decoded MEG signal occurred early in the trials, differences in heart rate between conditions manifested seconds later, during a time where the classifier no longer differentiated between approach-avoidance responses. This suggests that classification accuracy is not biased by the cardiac signal. Differences in eye movements are precluded by the counterbalanced nature of our design and could indeed not be used to predict threat/reward condition.

To summarize, this study shows that defensive freezing, indexed through cardiac deceleration, is associated with increased integration of reward and threat during approach-avoidance decision making. This threat-reward integration process and its moderation by defensive states is supported by low-frequency rhythmic activity in the dACC and is behaviorally relevant. These findings increase our understanding of how defensive responses such as freezing may facilitate adaptive threat-coping behaviors by upregulating the neural processing of potential reward and threat outcomes.

## Methods

This study was preregistered before data analysis on the Open Science Framework (https://osf.io/br9uz). All research activities were carried out in accordance with the Declaration of Helsinki, approved by the local ethics committee (Ethical Reviewing Board CMO/METC [Institutional Research Review Board] Arnhem-Nijmegen, CMO 2014/288), and conducted according to these guidelines and regulations.

### Participants

We recruited a total of 11 participants. One participant prematurely dropped out of data collection, and one additional participant was excluded due to irrational choice behavior (i.e., they approached threat-only trials with no reward / avoided reward-only trials with no threat more than 30% of the time – note that the M±SD approach/avoidance percentages on threat-only and reward-only trials for all other participants were 3±2.19 and 3.32±3.46, respectively). We thus included 9 participants for data analysis (completing 1 MRI and 5 MEG sessions each; aged 24±3 years, ranging 18-29 years, 5 females). Inclusion criteria were age (between 18 – 35 years old), English or Dutch speaking, and right-handedness, normal or corrected-to-normal vision. Exclusion criteria (based on self-report) included presence of metal in the body (such as a dental wire), pregnancy, epilepsy, claustrophobia, current or past treatment for major psychiatric disorders (such as anxiety, depression, mood disorders, and ADHD), cardiovascular conditions (such as heart rate arrhythmia), diagnosed low/high blood pressure, history of hormonal disorders or treatment, and MRI incompatibility (e.g., due to unremovable metal parts, active implants, or brain surgery). Participants were recruited through advertisements in student communication channels, and mainly consisted of research master students attending the Radboud University in Nijmegen, The Netherlands. All participants gave written informed consent prior to participation, were paid for participation (€105) plus bonus money contingent on their task choices (max €15 per session, see *Approach-avoidance conflict task* below).

### Experimental paradigm

#### Approach-avoidance conflict task

Participants performed a decision-making task in which they approached or avoided targets associated with varying reward and threat magnitudes (i.e., 0-5 euros/shocks; adapted from refs^8,17^; **Figure 1a**).

In each trial, after an intertrial interval of 8-10 s, participants were first presented with the cue screen detailing the money and shock levels on offer for that trial as stacks of orange and blue boxes (ranging 0-5 euro/shocks each, color-outcome relationships were counterbalanced between participants, the onscreen location of the orange and blue boxes was counterbalanced within participants, and colors were matched for lightness; i.e., HSL-values were [209, 59.2, 59.6] for blue and [25, 89.3, 59.6] for orange). More specifically, trials were separated in three trial types. First, in symmetrical trials (300 per participant), the objective money and shock levels were identical, such that equal information was presented on the screen between the two outcome types (i.e., money-shock levels were 1-1, 2-2, …, 5-5). Secondly, skewed trials (300 per participant) contained all remaining combinations of money and shock levels (i.e., 1-2, 1-3, 1-4, 1-5, …, 5-1, 5-2, 5-3, 5-4). Thirdly, in extreme trials (240 per participant), the level of one outcome was set to 0, while the level of the other was set to 5 (i.e., extreme reward = 5 euro, 0 shocks; extreme threat = 0 euro, 5 shocks). Thus, note that *only* in extreme trials one of the two outcomes could be set to a level of 0. After a fixed period of 1.5 s, the anticipation screen was presented for a variable interval of 1 – 4.5 s, containing the target (gray circle in the center) and the player icon (white square in the bottom). Because of the slow development of the heart rate signal over time, as preregistered, only trials with long intervals (i.e., ≥ 3.5 s, ±80% of all trials) were included in data analysis. The short trials were included in the design to ensure continued activation of the participant throughout the anticipation window (see e.g., refs^14,17^). After this interval, the target started to move either to the left or right side of the screen (counterbalanced), indicating the start of the response window. Throughout the duration of this target movement (800 ms) participants could either approach or avoid the target by positioning the player icon either on the same location as the target (approach) or on the opposite location (avoid) using buttons under their left and right index fingers. For example, if the target moved to the left, participants could jump to the left (using the left button) to approach the target, or jump to the right (using the right button) to avoid it. Approach choices were associated with a high probability of receiving either the specified money (40%) or shock (40%) level, and a low probability of no outcome (20%). Inversely, avoid choices were associated with a high probability of no outcome (80%) and a low probability of receiving the money or shock outcomes (10% each). If participants did not respond or responded too late, they would always receive the indicated number of shocks. The selected outcome was presented during an outcome screen following the response window (1.5 s). Shock outcomes were indicated by respective color coding of the target stimulus and delivery of the electrical shocks. Money outcomes were similarly indicated by color coding of the target stimulus, with an additional repeated presentation of a ‘+’ icon inside the target (number of presentations indicating the money level). In case of no outcome, the target simply remained gray, and in ‘missed’ trials the target was presented in red. The summed monetary outcome of three randomly selected trials per session (max. €15) was paid out as a bonus fee. The task was programmed in MATLAB^42^ using the PsychToolbox extension ^43^.

#### Procedure

Participants came to the lab for 1 MRI session (30 min.) and 5 MEG sessions (120 min. each). Upon arrival for the first (i.e., MRI) session, participants read and signed the informed consent and screening forms. Participants were then placed in the MRI scanner to collect 3 structural images: a standard T1-weighted anatomical image, a T2-weighted image, and a large-FOV (field of view) T1-weighted image optimized for imaging the participants’ skull. The last image was used to create a custom foam head cast to perfectly fit the participant inside the MEG scanner (see *head cast creation* below). Next, participants performed 5 MEG sessions. In each session, participants changed to MEG-compatible clothes (removing all metal objects) and were placed into the MEG scanner. After electrode placement and eye-tracker calibration (see *peripheral measurements and electrical stimulation* below), participants performed a standardized shock work-up procedure (as previously described by refs^17,44^). In this procedure, participants received and rated exactly five electrical shocks on a scale from 1 (not painful at all) to 5 (very painful). Shock intensities were divided in 10 levels (see *peripheral measurements and electrical stimulation* below) and were adjusted after each rating, aiming for a final shock intensity rating of 4 (i.e., uncomfortable but not painful; *mean±SD* of final shock intensity 4.57±1.34 out of 10). Next, participants read through the task instructions and performed 10 practice trials (without electrical shocks/monetary pay-out). In all following MEG sessions, participants were presented with a summary slide of the instructions rather than the full instructions (no practice trials). Participants then performed the approach-avoidance task, consisting of 3 runs of ±20 min. each per session. Although the individualized head casts (see below) greatly restricted head movements throughout the task, we measured participants’ head location using localization coils placed in both ear canals and on the nasion, and monitored this using online localization software^45^. In case of large deviations from the initial head position (>2 mm), we paused the experiment and instructed participants to move back to the original position. Participants’ head location in the first session was used as a reference for later sessions to minimize between-session variability in head location. At the end of the final session, we digitized participants’ head shape and the locations of the three localization coils using a Polhemus 3D tracking device, after which participants filled in a debriefing questionnaire. Monetary compensation was paid out after completing the final session.

#### MRI data acquisition

MRI scans were acquired using a Siemens MAGNETOM Skyra 3T MR scanner. A customized T1-weighted image (1 mm isotropic, TR = 4.5 s, TE = 1.59 ms, flip angle = 12, FOV = 256 × 256 × 208, interleaved order) was recorded for the purpose of head cast creation (see below). Another structural (anatomical) image was acquired (1 mm isotropic) using a T1-weighted 3D magnetization-prepared rapid gradient-echo sequence (MP-RAGE; TR = 2.3 s, TE = 3.03 ms, flip angle = 8, FOV = 256 × 256 × 192, ascending order). Finally, we recorded a T2-weighted image (fast low angle shot [FAST] sequence, TR = 4.5 ms, TE = 1.59 ms, slice thickness = 1 mm, FOV = 256 × 256 × 208) for other research purposes not reported here.

#### Head cast creation

Individualized head casts were created roughly based on the recipe provided by Troebinger and colleagues^40^. We extracted the head surface from the individual structural MRI using SPM12 (Statistical Parametric Mapping; Wellcome Trust Centre for Neuroimaging, London, UK) and used Fieldtrip to create a surface mesh of the head^46^. The head mesh was placed in the middle of a virtual copy of the MEG dewar using Solidworks 2022. After positioning, we added 3D objects to this head shape to ensure that the area in front of the eyes remains free of liquid foam, and clips to attach the head to a 3D-printed version of the MEG dewar. The 3D head shape was then sliced using PrusaSlicer software and printed using an Original Prusa i3 MK3S+ printer. After the head was fixated in a printed model of the dewar, FlexFoam-iT 15 (created by Smooth-On) was poured in the space between the printed head and DEWAR. After hardening, the head cast fit neatly around the participants’ head and into the MEG, minimizing movement and localization error. In contrast to Troebinger and colleagues, we did not include fiducial shapes into the cast, but used the standard ear and nasion placement to co-register and localize participants’ head position. This was done because pilots revealed that small movements were still possible with the head casts (mainly in vertical [z-axis] direction when relaxing) that we would not be able to correct for when fiducials were fixed to the cast.

#### Peripheral measurements and electrical stimulation

We measured respiration using a respiration belt placed around participants’ abdomen. Eye gaze and pupil diameter were collected using an Eyelink 1000 eye-tracker system (SR Research, Kanata, Ontario, Canada) at 1000 Hz (data not reported here). Electrical stimulation was performed using a Digitimer Constant Current Stimulator DS7A (www.digitimer.com) and standard Ag/AgCl electrodes. Shock intensities were divided into 10 levels, with the lowest and highest intensities stimulating at 1.9 mA / 6.9 mA respectively (both at 400 V) for a duration of 2000 μs (2 ms).

#### MEG data acquisition

MEG data were acquired using a whole-head CTF-275 system with axial gradiometers. Data were sampled at 1200 Hz after application of a 300 Hz low-pass filter. For four out of ten participants the following channels were permanently disabled due to high noise levels: MLC11, MLC32, MLC61, MLO33, MLT41, MRF66, MRO33, MRO52. For all participants, the helmet tilt was set at 50º and the viewing distance from the screen was about 80 cm. Stimuli were presented using a PROPixx beamer at a refresh rate of 120 Hz and a resolution of 1920 × 1080 pixels.

### Signal preprocessing

#### Magnetoencephalography

MEG data preprocessing and analyses were performed using MATLAB 2020b (The MathWorks), the Fieldtrip toolbox^46^, and custom analysis scripts. Data were epoched into trials ranging 1.5 s before trial onset to 2 s after the response window (7.5 s after trial onset). We performed third-order gradient field correction, baseline correction (demeaning), and applied a 200 Hz low-pass filter and a line noise (50/100/150 Hz) filter. Trials exceeding 1.5 mm of motion relative to the first trial were removed. Next, we performed manual trial rejection to remove trials with large deviations or artifacts, after which we used independent component analysis (ICA^47^) to remove noise components (e.g., heartbeat, eye blink). The heart rate components were saved for further analysis (see below). Finally, all trials were once more visually inspected to remove any remaining trials with large amounts of noise. We interpolated data for missing sensors (after removal due to high noise levels) using a weighted average of neighboring sensors. For classification analyses, the MEG data were further downsampled to 150 Hz.

#### Heart rate

As mentioned above, trial-by-trial timeseries of the cardiac signal were obtained by extracting them from the MEG signal using ICA^46,47^. To compute trial-by-trial heart rates, we performed peak detection on the extracted heart beat components and removed trials with poor peak detection (e.g., due to a noisy signal). Based on the heart beat peaks, we computed inter-beat intervals (IBIs) which were converted to heart rate in beats-per-minute (BPM = IBI/60). Heart rate timeseries were then baseline corrected relative to the mean heart rate during a pre-trial baseline window of 1 s (i.e., the baseline window is [−1, 0] relative to the cue screen onset). Trial-by-trial quantifications of heart rate (deceleration) were computed according to a ‘peak-to-trough’ approach. Based on the grand average heart rate timeseries across participants and conditions we selected the time point at which the average heart rate peaked (i.e., at around 3 s) and the latest time point before the start of the start of the response window (i.e., at 6 s). We then used these time points compute the trial-by-trial ‘slopes’ by subtracting the baseline-corrected heart rate at the ‘peak’ from the heart rate at the ‘trough’ (i.e., such that more negative slope values reflect stronger heart rate deceleration over time; *HR*_*slope*_ = *HR*_*trough*_ − *HR*_*peak*_).

### Data analyses

Generally, we used Bayesian Mixed-effects Models (BMMs) for all trial-by-trial analyses (*brms* and *rstan* packages in R and RStudio^48–50^). All models were estimated with 6000 samples (3000 warm-up) across 4 chains, followed a ‘maximal’ random-effects specification as recommended by Barr and colleagues^51^ (i.e., by-participant random intercepts, random slopes for all within-subjects variables, and all pairwise correlations between random intercepts and slopes). All models used the default weakly-regularizing priors provided by *brms*. Rhat values indicated that all models converged to a solution (i.e., all rhat values between 0.99 and 1.01).

Statistical inference was done on the highest density intervals (HDIs) of the posterior parameter distributions. Significance was determined based on the 95% and 90% HDIs, i.e., interpreted as ‘significant’ and ‘marginally significant’, respectively (see e.g., refs^52,53^ for more information on Bayesian indices of significance). Finally, we report effects’ posterior estimates (i.e., the mean values of the posterior distributions).

#### Behavior

Approach-avoidance choices were modelled using a Bernoulli distribution (logit link) and as a function of reward, threat, and heart rate deceleration as specified in equation (1):

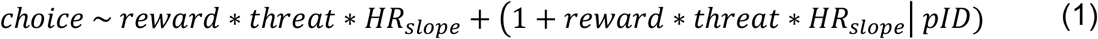

In this notation, the term left of the tilde reflect the dependent variable, whereas terms right of the tilde reflect the predictors (by-participant random effects in brackets). Asterisks indicate that the main effects as well as all interactions were estimated.

#### Heart rate

Trial-by-trial analyses of heart rate deceleration (as indexed by the slope) were performed according to the following model specification:

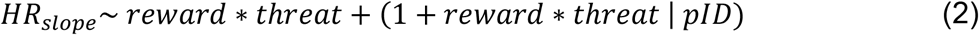

#### MEG classification analysis

*Classifier training*. All classification analyses were performed using the MVPA-light toolbox and its implementation in the FieldTrip toolbox in MATLAB^46,54^. For each participant, we trained a binary linear discriminant analysis (LDA) classifier on the extreme reward and extreme threat trials (max. 240 trials per participant). The classifier was constructed in a temporally-resolved way, such that it created a spatial filter that takes into account information of 15 consecutive samples (i.e., 100 ms). Training was done using k-fold cross-validation (10 folds, 2 repetitions) for every time point in a window ranging −1 to 6 s relative to the cue screen onset. For each participant, we selected the time point during the cue screen at which the classifier had the highest classification accuracy (computed as area under the curve, or ‘AUC’). These best-performing classifiers were subsequently used for cross-classification in the testing phase (see below).

#### Cross-classification

Best-performing classifiers per participant were used to cross-classify independent MEG data in the symmetrical and skewed trials by dragging the classifiers across the MEG timeseries per trial. Specifically, for each time point in each trial (taking into account the 100 ms/15 sample window around that timepoint), we let the classifier compute a value (TR value, ranging [−∞, +∞]) which essentially quantifies to what extent the MEG signal around that time point is more threat-like (positive TR values) or more reward-like (negative TR values). This way we created new trial-by-trial timeseries of decoded threat-reward information which was subsequently used for further statistical analysis.

#### Analysis of decoded threat-reward information

Trial-by-trial analysis of cross-classified TR values were performed by computing the mean TR value during the cue screen, anticipation screen, and their combination (per trial). These were subsequently analyzed (e.g., correlated to trial-by-trial choices and heart rate deceleration) using BMMs;

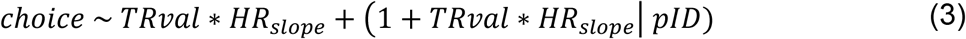

and;

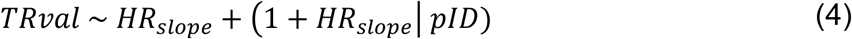

Trial-by-trial spectral analyses on threat-reward information during the cue screen were performed using Fourier transform.

#### MEG source reconstruction

We co-registered individual participant’s structural MRI scans to the MEG recording in two steps. First, we co-registered fiducial markers during the MEG acquisition to vitamin E capsules placed in ear-molds during the MRI session. This was followed by surface registration of the MRI-extracted head surface to a 3D head surface obtained with a Polhemus Fastrak device, using the Iterative Closest Point algorithm implemented in Fieldtrip. We then constructed a triangulated cortical surface mesh using the automatic surface extraction tool implemented in Freesurfer (recon-all: https://surfer.nmr.mgh.harvard.edu/fswiki/ReconAllTableStableV6.0). The resulting cortical meshes were registered to standard brains and downsampled to 7842 vertices using the HPC workbench software. The resulting mesh was used as source model. We used individual structural MRI images to create a single-shell head model for each participant, which was combined with the source model and the MEG gradiometer information to create a lead field. To reconstruct region of interest (ROI) activity (bilateral dACC and SMA), we first performed linearly constrained minimum norm beamformer based on the activity during the cue screen of all trials. The spatial filters for our four regions of interest were then applied to the MEG data and we extracted the dipole direction explaining most variance using principal component analyses. This was done for left and right dACC/SMA separately (four ROIs in total).

#### Coherence calculation

Because our experimental design is tailored towards maximum intra-subject reliability with many trials but few participants we opted to use trial-by-trial estimates of coherence. We computed coherence using a jack-knife procedure^27^. This procedure iteratively estimates the coherence based on all trials but one. The resulting coherence values are assigned to the left-out trial. This procedure highlights the contribution the left-out trial has on the overall coherence, whilst keeping the amount of data the same for each estimation. Note that the jackknife procedure reverts the direction of the coherence changes (i.e., lower values indicate stronger coherence).

Coherence between TR values (i.e., decoded threat-reward information) and ROI activity was computed in the window of cue presentation (i.e., 0 - 1.5 s relative to cue onset). We used Fourier transform with multi-tapering and a spectral smoothing of 5 Hz. Because we wanted to know to what extent neural signals in our ROIs were real-time correlated with the decoded TR-value timeseries, we focused on instantaneous (i.e., zero-lag) correlations by extracting the real-valued component of the complex-valued coherence coefficient.

Finally, to test how trial-by-trial coherence for four ROIs (left/right dACC/SMA) was related to cardiac deceleration, we performed a four-variable multivariate BMM (including the estimation of residual correlates between ROIs);

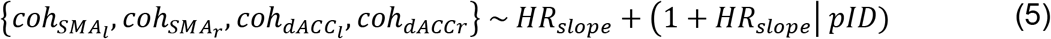

## Acknowledgements

This work was supported by a Consolidator Grant from the European Research Council (ERC_CoG – 2017_772337) awarded to Karin Roelofs, also supporting FK and LdV.

