## Supplemental Information for "Defensive freezing sharpens threat-reward information processing during approach-avoidance decision making"

### Classification of threat-reward information from eye movements does not exceed chance

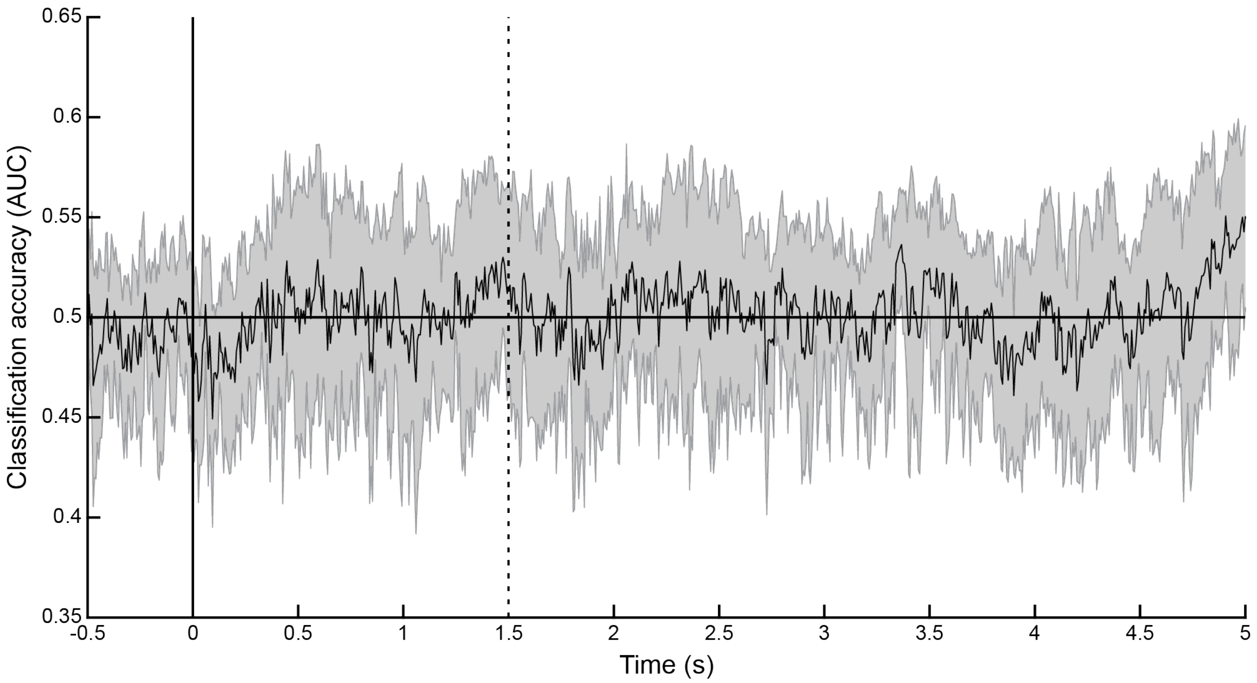


**Supplemental Figure S1. Classification accuracy of extreme threat and reward trials trained on eye movements.** *When threat-reward decoders were trained on eye movements instead of neural data recorded with MEG, classification accuracy (in AUC, area under the curve) of extreme threat and reward trials does not exceed chance (50%). Solid and dashed vertical lines represent cue and anticipation screen onset, respectively. Shaded area represents +/- 1 SEM.*

### Time-by-time generalization of threat-reward classifiers

To assess the temporal specificity of decoded threat-reward information from neural data, we investigated whether classifiers trained at each timepoint could generalize to other timepoints (time-by-time generalization, see e.g. ref[1]). The resulting generalization matrix revealed a pattern indicating that information was mostly specific within trial events. That is, classifiers trained during the cue screen performed well in other timepoints during the cue screen, but did not generalize to timepoints during the anticipation screen. This suggests that our classification picks up transients that reflect dynamics of the early decision[2], which might be qualitatively different from the information present in later timepoints. Inversely, classifiers trained at any point during the anticipation screen generally performed less well (i.e., had lower accuracy) but did generalize relatively well to other timepoints during the anticipation screen (**Supplemental Figure S1**).

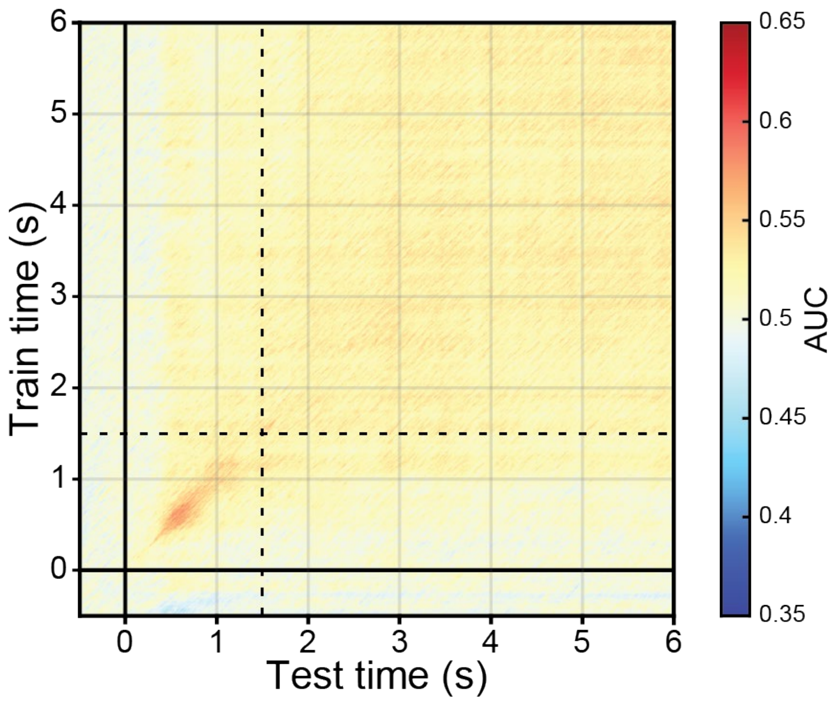


**Supplemental Figure S2. Time-by-time generalization of threat-reward classifiers.** *Classification accuracy (in AUC, area under the curve) of extreme threat and reward trials when tested (x-axis) and trained (y-axis) on various timepoints across the trial. Solid and dashed lines at t = 0 and t = 1.5 reflect cue and anticipation onset, respectively.*

### Correlating threat-reward information with trial-by-trial cardiac deceleration

There was no relation between cardiac deceleration and the average strength of threat-reward information, nor an interaction between average threat-reward information and heart rate on choice ($B_{HR}$ = 0.001, $HDI_{90\%}$ = [-0.002, 0.005]; $B_{TRval:HR}$ = 0.04, $HDI_{90\%}$ = [-0.11, 0.20]). This finding suggests that during freezing states neural information is not generally more threat-like or reward-like (averaged across time).

### Cardiac moderation of low-frequency coherence in the bilateral SMA

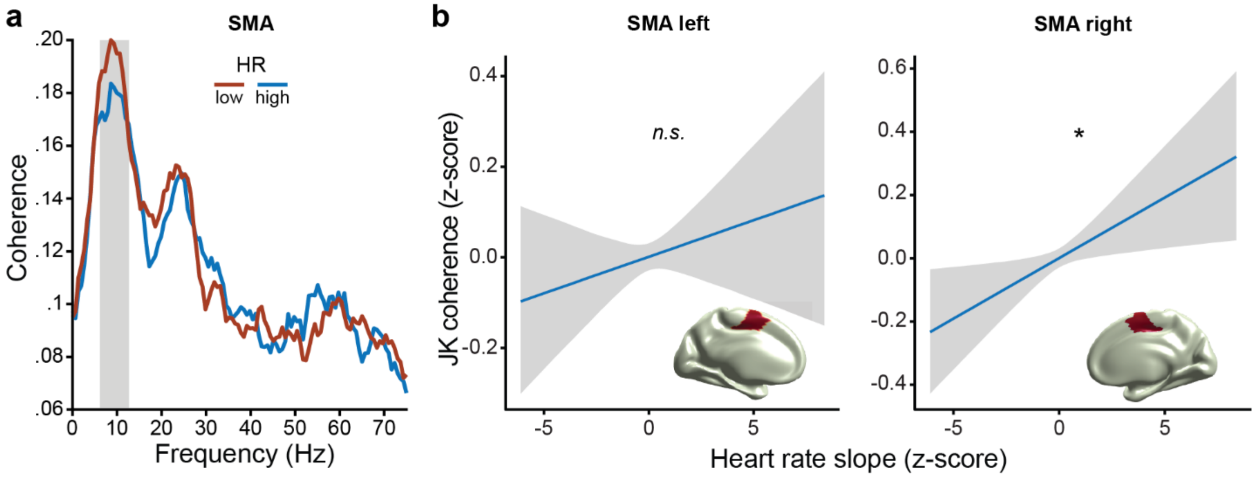


**Supplemental Figure S3. Cardiac moderation of low-frequency coherence in the bilateral SMA.** *(**a**) Coherence spectrum for reconstructed bilateral supplemental motor area (SMA) activity with decoded threat-reward information (averaged across hemispheres). For display purposes only, coherence is plotted separately for trials with 25% lowest vs. 25% highest heart rate (HR) slopes (i.e., both bins contain equal numbers of trials). Lower HR slope values indicate stronger cardiac deceleration. (**b**) Trial-by-trial jack-knife (JK) coherence in the left and right SMA with decoded threat-reward information in the 6-12 Hz range (indicated with gray shaded rectangle in **a**) is stronger during states of cardiac deceleration. Note that the y-axis is flipped; lower values indicate stronger coherence; similarly, lower heart rate slope values on the x-axis indicate stronger cardiac deceleration. Conditional-effects plots were extracted from the multivariate BMM. Asterisk indicates significant effect (i.e., $HDI_{95\%}$ does not include 0); ns = not significant (i.e., $HDI_{90\%}$ includes 0).*
